# MetaLogo: a heterogeneity-aware sequence logo generator and aligner

**DOI:** 10.1101/2021.08.12.456038

**Authors:** Yaowen Chen, Zhen He, Yahui Men, Guohua Dong, Shuofeng Hu, Xiaomin Ying

## Abstract

Sequence logos are used to visually display conservations and variations in short sequences. They can indicate the fixed patterns or conserved motifs in a batch of DNA or protein sequences. However, most of the popular sequence logo generators are based on the assumption that all the input sequences are from the same homologous group, which will lead to an overlook of the heterogeneity among the sequences during the sequence logo making process. Heterogeneous groups of sequences may represent clades of different evolutionary origins, or genes families with different functions. Therefore, it is essential to divide the sequences into different phylogenetic or functional groups to reveal their specific sequence motifs and conservation patterns. To solve these problems, we developed MetaLogo, which can automatically cluster the input sequences after multiple sequence alignment and phylogenetic tree construction, and then output sequence logos for multiple groups and aligned them in one figure. User-defined grouping is also supported by MetaLogo to allow users to investigate functional motifs in a more delicate and dynamic perspective. MetaLogo can highlight both the homologous and nonhomologous sites among sequences. MetaLogo can also be used to annotate the evolutionary positions and gene functions of unknown sequences, together with their local sequence characteristics. We provide users a public MetaLogo web server (http://metalogo.omicsnet.org), a standalone Python package (https://github.com/labomics/MetaLogo), and also a built-in web server available for local deployment. Using MetaLogo, users can draw informative, customized and publishable sequence logos without any programming experience to present and investigate new knowledge on specific sequence sets.

## Introduction

Sequence logo was first proposed by Schneider and Stephens in 1990 [1], and has been widely used for sequence pattern visualization in academic fields. Each position of a sequence logo is stacked by different amino acids or nucleotides, with the height of each base indicating its degree of conservation at that position. The most commonly used sequence logo generators include Weblogo [2], Seq2Logo [3], ggseqlogo [4], Logomaker [5], RaacLogo [6] and others, involving web servers, Python and R packages, etc. By showing graphical representations of sequences, these tools greatly accelerate researchers’ exploration of sequence patterns and motifs.

However, a common problem is that most sequence logo tools were designed to highlight the conservations among sequences and to underrate the variations. The more diverse the positions, the less signal can be indicated in sequence logos. This setting could help to reveal the real conserved motifs if the input sequences are indeed homologous. But when there exists intrinsic divergence among sequences, it is difficult for conventional sequence logo generators to sort out clues. In other words, ortholog and paralog of sequences need to be considered when performing sequence analysis, especially for motif discovery. Among the sequenced data of natural DNA or amino acids, there may exist artificial synthesis and contaminated sequences, which also need to be recognized when drawing sequence logos. Take the B cell receptor (BCR) as an example. Complementarity determining region 3 (CDR3) is the most hypervariable region in BCR. It is known that BCRs binding to different targets may have different CDR3 sequence characteristics, involving length, motif, polarity, physical chemical properties, structures and so on. Researchers often utilized sequence logos to reveal functional motifs of antibodies with specific affinity towards certain antigens. But experiment-selected sequences always contain containments, false positive ones or multi-specific antibodies which also bind to other targets. If all the sequences are taken as input for sequence logo generating, BCRs with different specificities are mixed and the functional motifs could be covered by interference signals, which makes the sequence analysis challenging. Besides CDR3s analysis, other motif-related studies, such as transcript factor motif analysis, clustered regularly interspaced short palindromic repeats (CRISPR) array analysis, evolutionarily conserved sequences analysis and others, all have the same demands. Therefore, researchers need a convenient tool which could sort out the heterogeneity of the data and display pattern dynamics across different clusters with specific functions, so as to understand the sequence characteristics in a more delicate manner.

To solve the problems mentioned above, we developed MetaLogo, which take heterogeneous sequences as input, then perform sequence clustering based on the phylogenetic tree or user-defined grouping, and finally output sequence logos for each group and place them aligned in one figure. MetaLogo can highlight both the conserved motifs and diverse regions among groups, and can possibly relate these homogeneities and heterogeneities to gene functions. Since the grouping are based on the phylogeny, MetaLogo could also be used to infer functions or taxonomies of target sequences from their neighbors in the same group. Since MetaLogo does not assume that the input sequences are from the same homologous group, the motifs indicated by sequence logos from MetaLogo could be more informative and reliable than those from conventional sequence generators. To use homology information beyond the input sequences, users can use Consurf [7] and MetaLogo together to get all the sequences of the homology group into MetaLogo and construct the sequence logo diagrams.

## Description

MetaLogo provides a public web server (locally deployable), and a stand-alone Python package at the same time to provide researchers with the most convenient service. Users can input files in *Fasta* or *Fastq* format, and specify the grouping strategy to build sequence logos. Two grouping strategies are supported, including auto-detection grouping and user-defined grouping.

### Grouping strategies

The pipeline for auto-detection grouping scenario is showed in Figure 1. MetaLogo firstly deduplicates the input sequences to reduce the computational cost. Then multiple sequence alignment (MSA) and phylogenetic tree construction are performed sequentially. MSA is conducted by Clustal Omega [8], and phylogenetic tree building is conducted by FastTree 2 [9]. TreeCluster [10] is utilized for sequence clustering based on the phylogenetic tree. Supported parameters for clustering methods includes ‘max’, ‘max_clade’ and ‘single_linkage’. Since a threshold is needed for distance-based clustering, MetaLogo introduces a resolution value (from 0 to 1) to control the clustering process. MetaLogo first calculates pairwise distances among all the nodes of the tree, then it sorts the distances and maps the indices range of the sorted distances to range [0, 1], and finally choose the index and the corresponding distance as the threshold according to the user-provided resolution.

**Figure 1.**
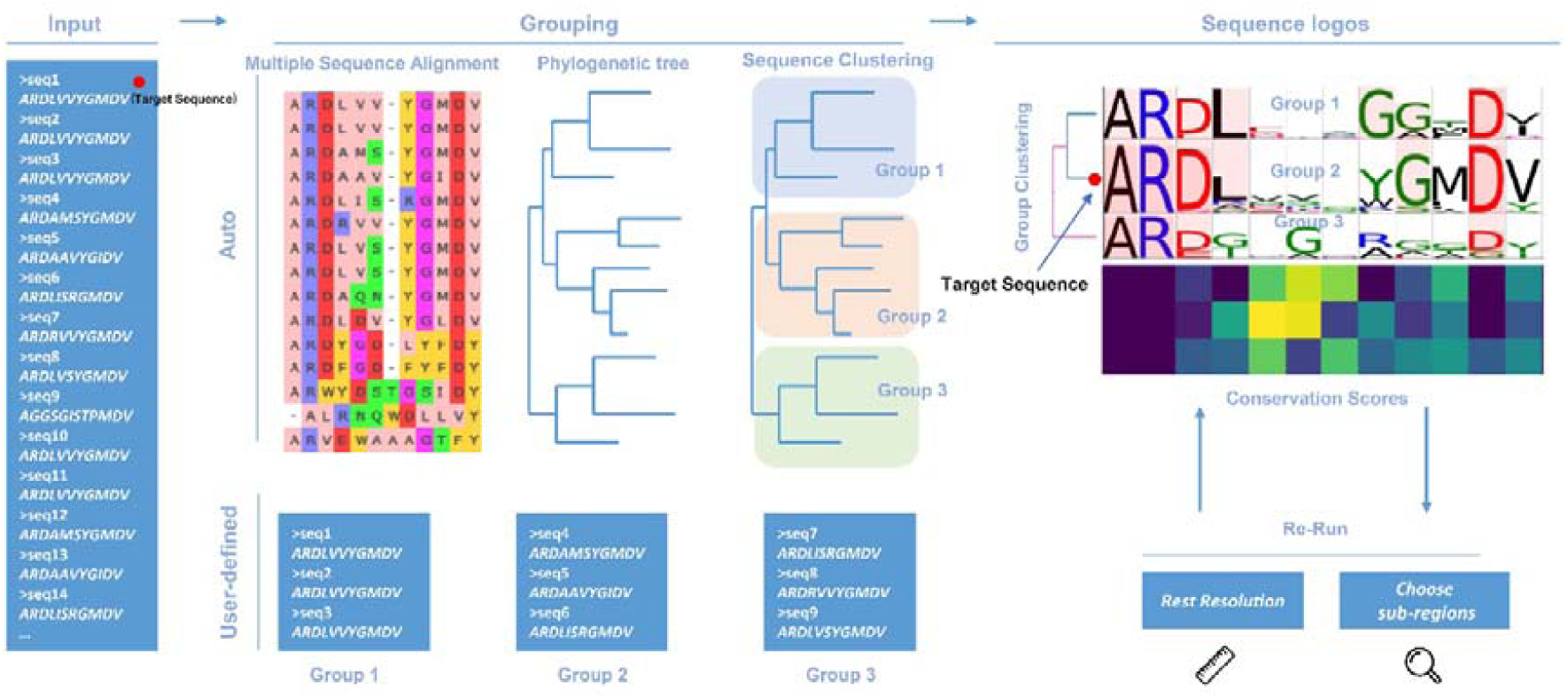
The schematic diagram of sequence logo generating process by MetaLogo. Sequences can be automatically clustered based on evolutionary relationships, or they can be grouped according to user-specified rules. MetaLogo draws and aligns sequence logos for each group, and reports the conservative status of each groups. Users can adjust the cluster resolution and sequence subregions to redraw sequence logos. The first sequence of the input will be tracked in its position in the final grouping.

When the resolution is set to 1, MetaLogo will treat all the sequences as a whole homologous group and output a single sequence logo for it. When the resolution is set to lower values, more groups could be identified based on the evolutionary relationships among the sequences. The MetaLogo public server provides a fast re-run module for users to quickly modify the resolution value to give them a dynamic view on the data.

In the auto-detection grouping mode, MetaLogo could also be used to annotate the target sequence. Take the target sequence together with other sequences with known annotations, MetaLogo could indicate the position of the target sequence in the phylogenetic tree and infer its functions according to its neighbors. The group specific motif showed by the sequence logo could also suggest the important functional sites of the target sequence. The MetaLogo public server treats the first sequence of the input as the target sequence and uses a red dot on the output figure to represent its location (Figure 1).

Users can also specify their own grouping rules. A simple one is to group sequences by sequence lengths, which may not make sense from an evolutionary perspective, but might be needed under certain scenarios. Users can also group their sequences in advance, and input sequences to MetaLogo with grouping information indicated in the sequence names. In this mode, MetaLogo also tries to align sequence logos by an adjusted MSA algorithm, which will be introduced in the following part.

### Sequence logo representation and similarity

For each set of sequences, MetaLogo first calculates the information contents of amino acids or nucleotides at each position in bits [11]. In order to align different sequence logos, to identify the consistently conserved motifs among groups, and also to cluster the sequence logos, we need to measure the similarities between bit arrays of positions from different logos. For example, *P* and *Q* are bit arrays of positions from two different protein logos and defined as follows:

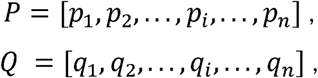

where *n* is the number of amino acids types and item *p*_*i*_ and*q*_*i*_ represent the information contents of the *i*th amino acid in the two positions, specifically. The arrays are sorted based on a fixed amino acids order.

To measure the similarity between *P* and *Q*, MetaLogo provides dot production (DP) and cosine similarity (COS) for users to choose from, which are commonly used as similarity measures and defined as follows:

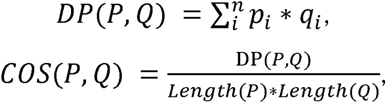

where *Length*(*P*)and *Length*(*Q*) represent the length of vector *P* and *Q*.

Besides bit arrays, we could also use frequency arrays to measure the similarity between positions. For each amino acid in one position, its frequency could be treated as the probability of one sequence having it in that position. Thus, here we could use similarity measurements designed for probability distributions.

MetaLogo allows users to choose the Jensen–Shannon divergence (JSD; [12]) as the similarity measurement. The JSD is a method of measuring the similarity between two probability distributions, and is a symmetrized version of the Kullback–Leibler (KL) divergence [13]. Note in the following context, *P* and *Q* represent discrete probability distributions which sum to one. JSD is defined as follows:

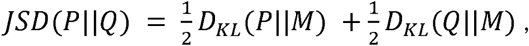

where 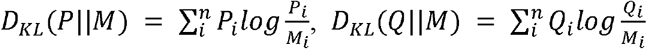 and 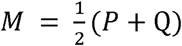.

Bhattacharyya coefficient (BC) [14] could also be used as a similarity measurement for two statistical samples. Since probability array does not indicate conservation like bit array do, hence MetaLogo provides an entropy (H) [15] adjusted Bhattacharyya coefficient (EBC) as a choice to measure the probability array similarity, which is defined as follows:

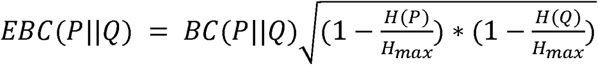

where 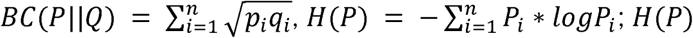 is the entropy of *P* and *H*_*max*_ is the max entropy for a *n*-dimensional probability vector.

Among these measurements, COS and JSD consider both nonconservative and conservative patterns, whereas DP and EBC only value conservative patterns among groups.

### Sequence logo alignment and clustering

In the user-defined grouping scenario, MetaLogo performs sequence logo alignment to indicate the dynamic changes of motifs among groups. The alignment between sequence logos is based on the Needleman–Wunsch algorithm, which is a classic global sequence alignment algorithm. For global multi-logo alignment, MetaLogo adopts the method of progressive alignment construction [16]. The closest pair of sequence logos are aligned first, and the next logo closest to the aligned sequence logo set is successively added for alignment. Introduced gaps and inserts of each alignment are retained for subsequent alignments until all logos get aligned. Users need to specify a certain similarity metric we mentioned above, and also the penalty for inserts and gaps.

In both auto-grouping and user-defined group scenario, MetaLogo can visually highlight the highly similar pairs of positions between groups by connecting them using colorful strips.

After sequence logo are generated for each group, MetaLogo clusters the groups to reveals the correlations and distances among groups, which needs to be consistent with the phylogenetic tree of all the sequences. Representative vectors of each group are submitted to a hierarchical clustering process with the UPGMA (unweighted pair group method with arithmetic mean) method. A resulted dendrogram is added to the sequence logos to show the group relations.

### Visual capacity for long sequences and large amount of sequences

For longish sequences, MetaLogo provides parameters for users to specify the focused range of the alignment to make a clear organization of the plot. For large amount of sequences, many unexpected groups could be identified. Since it is not easy to present them all on one figure. MetaLogo allows users to choose the top *N* groups with the most sequences, where *N* is the number of groups users want to keep on the figure.

### Layouts and styles

MetaLogo supports four different logo layouts, including horizontal, circular, radial and 3D layouts. These diverse layouts are suitable for different scenes specifically. The horizontal layout is the default one, which can deal with most scenarios; the circular layout can more clearly show the conservations across multiple sequence groups; the radial layout is suitable to display sequences with conservative motifs in the middle or at the end of the sequences, rather than at the beginning; the 3D layout makes sequence logos more diverse and aesthetic.

MetaLogo allows customization of most of the operable elements in the figure, including figure size, ticks size, label size, labels, title, grids, margins between items, colors of items and so on. Users can also choose whether to display axis, ticks, labels, group ids, etc. Multiple formats of figures are supported, including *PNG, PDF, SVG, PS* and *EPS*.

### Basic statistics and intermediate results

MetaLogo provides users basic statistics of their input data to help users get a more complete understanding of the data. An entropy heatmap is utilized to indicate the conservation of each position in each group (Figure 1). A boxplot is used to display the entropy distribution of each group. A clustered heatmap is also provided to show correlations between every pair of groups, which could help users to understand which groups are more similar in conservation pattern and which groups are more divergent. Distribution of pairwise distances of sequences in the phylogenetic tree is also provided by MetaLogo, which could help users to adjust the clustering resolution value.

Most intermediate results during the sequence grouping and logo generating are provided by MetaLogo for visualization and downloading. Users can view the MSA result and phylogenetic tree on the website and can also download them for further analysis.

### Package Install and server deployment

Users can directly access our public web server (http://metalogo.omicsnet.org), or install MetaLogo as a Python package locally. Three examples of sequence sets are provided with the codes and on the server. One set contains sequences of *Escherichia coli* transcription factor binding sites [17], the second set contains sequences of CDR3s of verified antibodies detected in BCR repertoires of individuals with COVID-19 (coronavirus disease 2019; [18]), the third set contains sequences of another experiment validated BCR clonotypes from patient with COVID-19 [19]. A detailed tutorial for MetaLogo is provided online (https://github.com/labomics/MetaLogo/wiki). After the installation, users can run MetaLogo directly in the system terminal or import MetaLogo functions into their own scripts or projects. Users can also deploy MetaLogo as a web service on their local area network through Docker, which is convenient for people with no programming experience. Relevant parameters could be set for MetaLogo web server, including the number limitation of allowed sequences and the size limitation of uploaded files, etc.

## Use cases

### Case 1: clonotypes of BCRs from COVID-19 patients

In the past 2 years, the COVID-19 pandemic has brought obvious impact on human’s health and social activities all over the world. Many laboratories have been seeking clues from blood of COVID-19 patients to help treat the disease, and several potential therapeutic antibodies have been discovered [20]. Among these exploratory studies, Galson *et al*. deeply sequenced BCRs from a cohort of COVID-19 patients and discovered convergent immune signatures towards severe acute respiratory syndrome coronavirus 2 (SARS-CoV-2; [19]). In total 463 convergent clonotypes, or representative CDR3s, were detected, after matching to earlier published studies. These clonotypes were detected in at least four of the COVID-19 patients, but not present in healthy controls or individuals following seasonal influenza vaccination, which means they might response specifically to SARS-CoV-2. Since they did not focus on motifs of these sequences, we downloaded the CDR3 sequences of these clonotypes as well as immunoglobulin heavy chain variable region gene (IGHV) and immunoglobulin heavy chain joining gene (IGHJ) gene annotations and tried to use MetaLogo to reveal the conservation patterns among these sequences.

Figure 2A shows the multi-group sequence logo of all the 463 clonotypes. MetaLogo automatically divided the sequences into different groups according to their evolutionary relationships with a given resolution of 0.8. Conserved motifs shared by adjacent groups are connected by light blue bands. It is obvious that the heads and tails of all the groups show conserved patterns, whereas the middle of them seem more diverse. It is worth noting that there is an obvious GSY motif in the middle of the sequences in the groups 11 and 12, and a YYYGM motif near the tail of the group 4. Figure 2B shows the sequence counts of each group; the group 4, 11 and 12 account for >50% of the total sequences. Figure 2C shows entropies of each position of each group. The higher the entropy, the less convergent the position. Figure 2D shows boxplots of the entropies of each group and the group 11 shows the most conserved sequence pattern with the lowest median entropies. Figure 2E shows the clustering result of the groups and it seems that the group 4 is the most different one from others. Since the groups 11,12 and 4 attract our attention, we checked the V and J gene annotations of sequences in these groups. Interestingly, most sequences are from IGHV 3-30 and IGHV 3-30-3 (Figure 2F). IGHV 1-24 is also involved in annotations for sequences of the group 12. As to J gene, sequences in the group 4 are almost all from gene IGHJ 6, which provides the YYYGM motif for the sequences (Figure 2G).

**Figure 2.**
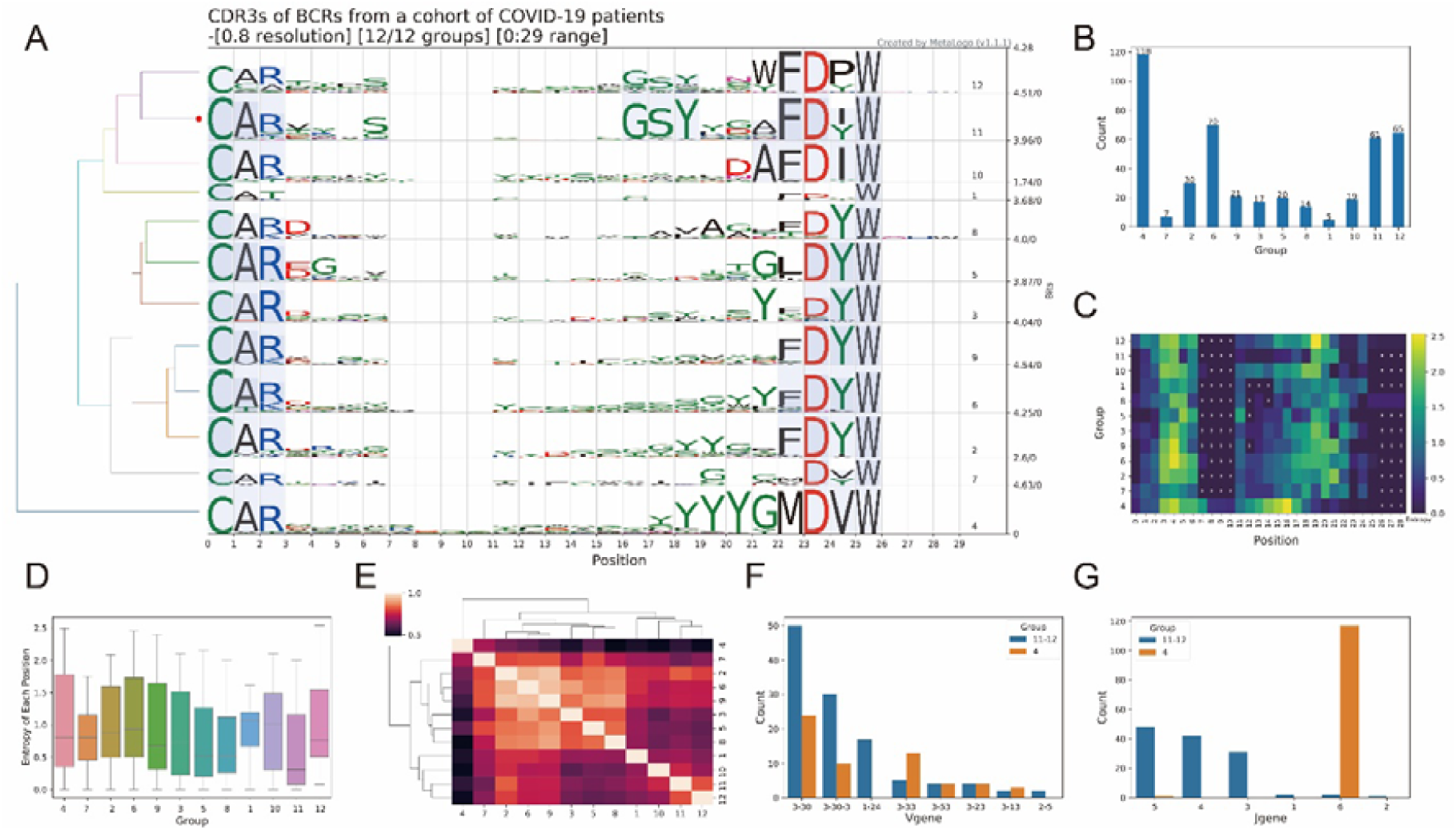
Sequence logos and analysis results from MetaLogo on the BCR clonotypes. **A**. Sequence logos for all the BCR clonotypes. The left tree indicates the relationships among groups. The red dot on the tree indicates the group to which the target sequence (first sequence of the input) belongs. Conserved positions among groups are connected by colored strips. **B**. Sequence counts of each group. **C**. Entropy heatmap of each group. ‘X’s represent gaps. **D**. Boxplot of entropies of positions in all groups. **E**. Clustering result of sequence logo groups to reveal the relationships among them. **F**. The V gene annotation distribution for the group 11-12 (group 11 and group 12) and the group 4. **G**. The J gene annotation distribution for the group 11-12 (group 11 and group 12) and the group 4.

According to the above results from MetaLogo, we can conclude that GSY and YYYGM are two potential motifs in the CDR3 region of BCRs from patients with COVID-19. These two motifs are mostly contributed by IGHV 3-30 (or 3-30-3) and IGHJ 6, respectively. We checked the published potential antibodies against SARS-CoV-2 from OAS (Observed Antibody Space) [21] database and found many antibodies contain the motif GSY and YYYGM in their CDR3 sequences, which is consist with our results but reminds the necessity of in-depth study of this phenomenon. Figure 2A-E were all generated by MetaLogo and the results indicate the capacity of MetaLogo to discover valuable knowledge among sequences in a convenient way.

### Case 2: coronavirus spike glycoprotein S1, C-terminal

To further demonstrate the function of MetaLogo to annotate sequences with unknown functions or evolutionary positions, we downloaded all the sequences of the CoV_S1_C family (PF19209) from Pfam [22]. The CoV_S1_C domain family contains sequences from the C-terminal of coronavirus spike glycoprotein S1, which could be related to the viral entry process into the host cell. In total 72 sequences were downloaded from the family. Figure 3A shows the sequence logos generated by MetaLogo with an auto-grouping resolution of 0.2. MetaLogo automatically considers the first sequence of the input as the target for annotation. MetaLogo indicates that the target sequence is assigned into the group 5 (red dot on Figure 3A). All the sequences are grouped into six clusters based on the phylogenetic tree given a resolution of 0.2. The clustering of representation vectors of these sequence logos (Figure 3B) and the pairwise distance distribution of the sequences (Figure 3C) indicate that these groups could be divided into two main parts. We investigated the taxonomies of these sequences and found that the group 3, 2 and 5 are classified as *Betacoronavirus*, the group 10, 9 are classified as *Alphacoronavirus*, whereas the transitional groups 6 and 12 contain sequences from *Alphacoronavirus, Betacoronavirus* and *Deltacoronavirus* (Figure 3D). The result makes sense from an evolutionary perspective.

**Figure 3.**
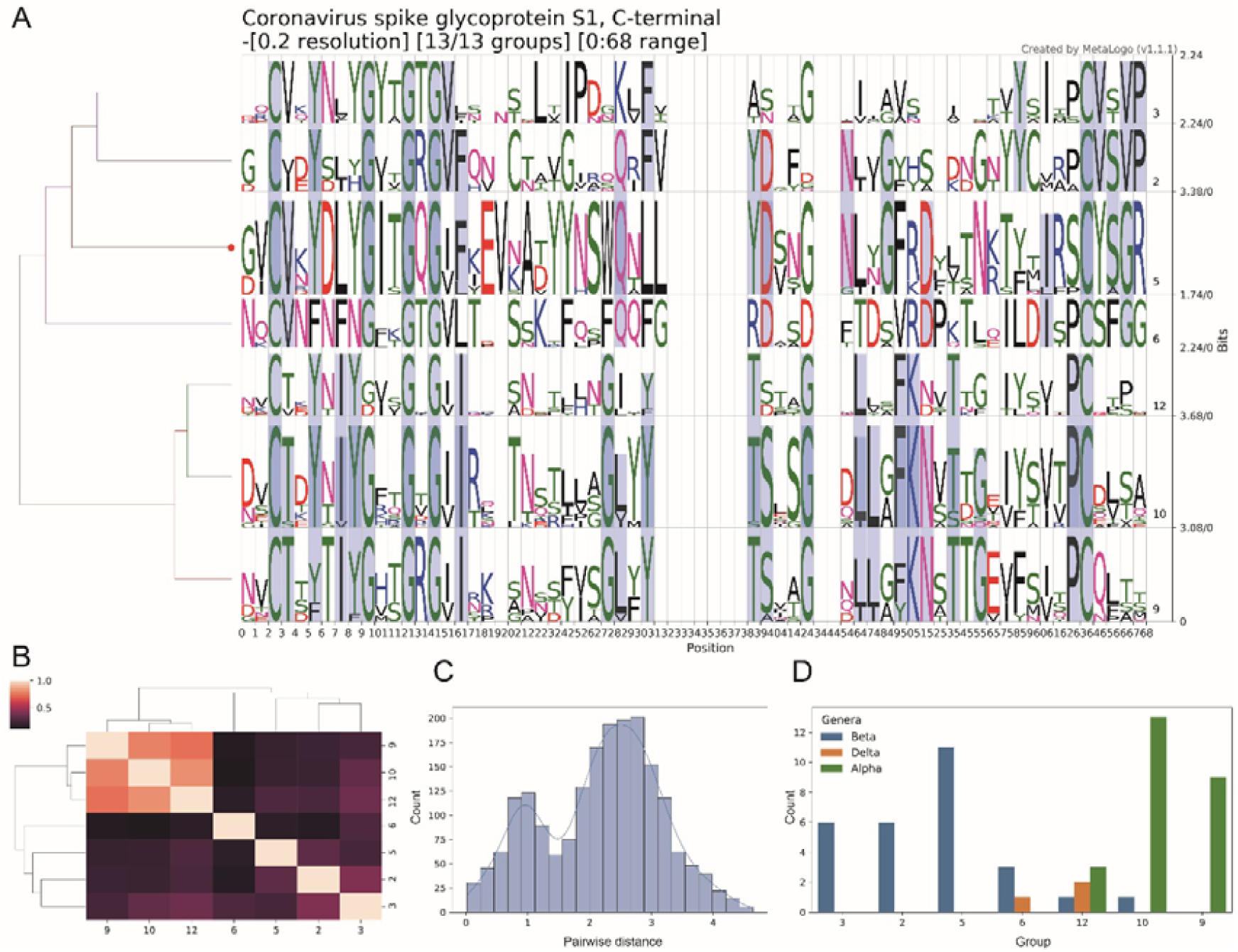
Sequence logos and analysis results from MetaLogo on the C-terminal sequences of the *Coronavirus* spike S1 protein. **A**. Sequence logos for all the S1 C-terminal sequences. The left tree indicates the relationships among groups. The red dot on the tree indicates the group to which the target sequence (first sequence of the input) belongs. Conserved positions among groups are connected by colored bands. **B**. Clustering result of sequence logo groups to reveal the relationships among them. **C**. Distribution of pairwise distances of the nodes in the phylogenetic tree. **D**. Taxonomy classifications of sequences in each group.

From the results, we can conclude that our target sequence is likely from *Betacoronavirus*, specifically *Embecovirus*. Meanwhile, from the results we can tell the motifs of the C-terminal domain of the S1 protein in *Embecovirus*, such as the *YYNSWQ* motif and the *IRSCYSGR* motif. Corresponding sequences from *Alphacoronavirus* are more conserved than those from other genus, especially *Betacoronavirus*. Further investigations are needed to study the relationships between the motifs and the protein functions. Figure 3 A-C were generated by MetaLogo. These results show that MetaLogo could perform reliable investigation on the sequence population and also support informative annotation for a single sequence.

## Conclusion

MetaLogo is a new generator for customized, informative and publishable sequence logos. Unlike existing tools, MetaLogo can automatically infer the evolutionary groups among the sequences and draw multiple sequence logos for different groups in one figure. MetaLogo is good at uncovering homogeneity as well as heterogeneity among sequences. User-defined grouping and statistical analysis are also supported by MetaLogo for users to reveal the pattern dynamics across groups. MetaLogo provides a free web server for public use, as well as a stand-alone Python package and a docker web service for local deployment. We will value the suggestions and comments from users, and continue to maintain code updates and upgrades to continuously contribute to the community.

## Key Points

- MetaLogo is a new sequence logo generator aware of heterogeneity of the sequences, it can draw sequence logo for each evolutionary group considering the phylogenetic relationships.
- For user-defined grouping, MetaLogo can perform pairwise and global sequence logos alignment to highlight the sequence pattern dynamics across different sequence groups.
- MetaLogo provides basic statistical analysis to additionally reveal the sequence convergences and divergences among sequence groups.
- MetaLogo provides a public web server, a deployable local web server with docker, as well as a stand-alone Python package for making highly customized sequence logos.

## Acknowledgments

We thank our colleagues in the lab including Pu Liu, Chao Feng, Sijing An and Runyan Liu for their careful reviews and feedbacks on the MetaLogo web server.

